# 5-hydroxy-4′-nitro-7-propionyloxy-genistein inhibited invasion and metastasis via inactivating wnt/β-catenin signal pathway in human endometrial carcinoma JEC cells

**DOI:** 10.1101/250605

**Authors:** Jun Bai, Binjian Yang, Xin Luo

## Abstract

Chemotherapy has been demonstrating more important roles in the treatment of carcinoma, but drug resistance and side effects restrict its usage in clinic, so developing new type of drug with low side effects and low-drug resistance has become a hot researching spot. The present study aimed to investigate the anticancer effects of 5-hydroxy-4’-nitro-7-propionyloxy-genistein (HNPG) and elucidate its underlying molecular mechanism. The inhibitory effects of cell viability of HNPG were detected using MTT assay, flat plate clone formation method and Transwell assay. The distribution of cell cycle was analyzed using FCM method. The morphological alteration, root-mean-squared roughness (Rq), average roughness (Ra), Young's modulus and adhesive force were measured by AFM. qRT-PCR and western blotting analysis were used to explore the possible molecular mechanism. It was found that HNPG presented with dramatic activity against JEC cells *in vitro*, inhibited the proliferation and colony formation, attenuated invasion and migration ability, arrested cell cycle in G1 phase in dose-dependent manner. Simultaneously, cell body shrunk and pseudopod structure retracted, Rq and Ra reduced, Young's modulus and adhesive force increased, accompanied by β-catenin, C-myc, Cyclin D1, MMP-2, MMP-7 and MMP-9 down-regulated. In summary, HNPG may be a novel candidate for chemotherapeutic drug.

## INTRODUCTION

Endometrial carcinoma (EC) is a group of epithelial malignancies that occur in the endometrium, and endometrial adenocarcinoma is the most common form accounting for 80 % ~ 90 % of all pathological types (Ethier et al., 2017). EC is one of the three malignant tumors of female genital tract, accounting for 7 % of malignant tumor in human carcinomas, and 20 % ~ 30 % of female malignant tumors (Kim et al., 2017). The average age suffering EC is 60 years with 75 % of patients aged 50 years or older. The morbidity of EC has been increasing worldwide in recent years (Ding et al., 2017), the main treatment of EC is surgery, radiotherapy, chemotherapy and hormone therapy (Ethier et al., 2017; Kim et al., 2017). Chemotherapy is one of the most important treatment measures for the EC of late-stage and recurrence, and it can also be used to EC with high-risk factors of recurrence (Kim et al., 2017; Ding et al., 2017). Nowadays, chemotherapy drugs, such as cisplatin, paclitaxel, cyclophosphamide, fluorouracil, mitomycin, etoposide and so on, are commonly used in clinical chemotherapy (Lester-Coll et al., 2017). Although chemotherapy demonstrates more and more important role in the treatment of EC, the generation of drug resistance and its side effects restrict its usage in clinic. Therefore, developing new type of drug with low side effects or without side effects and low-drug resistance has become a hot researching spot.

Genistein (GEN), one kind of isoflavone extracted from soybean, is reported that possess extensive antitumor activities via regulating oncogenes in many signal pathways, such as anti-proliferation, inhibitive metastasis, induction of apoptosis and so on (Zhong et al., 2017). However, the lower absorption rate of gastrointestinal tract and lower biological activity limit its clinical application (Zhong et al., 2017). It is well known that structural modification could enhance the solubility and biological activity (Chen et al., 2017), 5-hydroxy-4’-nitro-7-propionyloxy-genistein (HNPG), a novel synthetic derivative of GEN, possesses a nitro group in C-4’, a hydroxyl group at C-5 and a propionyloxy group at C-7 (Wang et al., 2012; Wang et al., 2005), which mean HNPG possess more solubility and biological activity than its precursor GEN. It was reported that HNPG exhibited an inhibition of proliferation in gastric and breast cancer, but its antitumor effect has not been examined in EC, and its molecular biological mechanism has not been investigated (Wang et al., 2012; Wang et al., 2005).

Wnt/β-catenin signal pathway is a complex and conservative signal pathway in mammal and human, is composed of a series of interacting proteins, regulates cell proliferation, adhesion, movement and so on, plays an important role in embryo development and tissue repair (Krishnamurthy and Kurzrock, 2018; Shanmugapriya et al., 2017). Abnormal activation of wnt/β-catenin signal pathway can cause abnormal proliferation and differentiation and result in tumor occurrence (Krishnamurthy et al., 2018). In addition, wnt/β-catenin signal pathway may induce the process of epithelial mesenchymal transformation (EMT), and affect the progression and metastasis of carcinoma (Shanmugapriya et al., 2017). So far, researches have shown that wnt/β-catenin signal pathway plays an important role in the occurrence of EC, and about 40 % of EC exhibit the abnormal activation of wnt/β-catenin signal pathway, especially in endometrial adenocarcinoma (EEC) (Eritja et al., 2017). Researchers discovered that the mutation of CT-NNB1 exon-3 could cause β-catenin protein accumulation, activated wnt/β-catenin signal pathways via analyzing 271 cases of EEC patients (Liu et al., 2014). It was reported that the expression of wnt10b was significantly increased in the tissues of EC (Chen et al., 2013). Researchers illuminated the role of a new signal pathway of PCDH10-wnt/β-catenin-MALAT1 on occurrence and progress in EC (Zhao et al., 2014).

In the present study, the anticancer effects of HNPG in JEC cells were evaluated *in vitro*, and the results demonstrated that HNPG could inhibit proliferation, clone formation, invasion and metastasis, which might attribute to HNPG increasing the cell accumulation in G1/S phase, altering cell morphological features, decreasing Rq and Ra, enhancing Young's modulus and adhesive force in JEC cells through inactivating wnt/β-catenin signal pathway, down-regulating β-catenin, C-myc, Cyclin D1, MMP-2, MMP-7 and MMP-9, which not only demonstrated the antitumor effects of HNPG, but also highlighted the basic molecular biological mechanism of HNPG compared with those previous reports (Wang et al., 2012; Wang et al., 2005).

## RESULTS

### HNPG inhibited the proliferation of JEC cells

JEC cells were treated with different concentrations of TAX, GEN and HNPG respectively ranging from 0.01~12.8 μM, 0.5~128 μM and 0.125~32 μM for 24 h, and the proliferation of JEC cells was inhibited by TAX, GEN and HNPG in a dose-dependent manner and IC50 were 0.78 μM, 16.24 μM and 3.86 μM respectively. TAX of 0.8 μM, GEN of 16μM, HNPG (2, 4 and 8 μM) were used to subsequent experiments to measure the inhibitive rates for 24, 48 and 72 h. The proliferation of JEC cells was markedly inhibited in a dose-and time-dependent manner, the inhibition rate of every HNPG-treated group was significantly different compared with the control group (P_2μM/0.1%DMSO_<0.05, P_4μM/0.1%DMSO_<0.05, P_8μM/0.1% DMSO_<0.05), and there was statistical difference among each HNPG-treated group (P_2/4μM_<0.05, P_2/8μM_<0.05, P_4/8μM_<0.05), and there were no statistical difference among HNPG of 4 μM, GEN of 16 μM, and TAX of 0.8 μM (P_HNPG of 4μM/GEN of 16μM_>0.05, P_HNPG of 4μM/TAX of 0.8 μM_>0.05, P_TAX of 0.8 μM/GEN of 16μM_>0.05) (Fig. 2 A-D).

**Figure 1.**
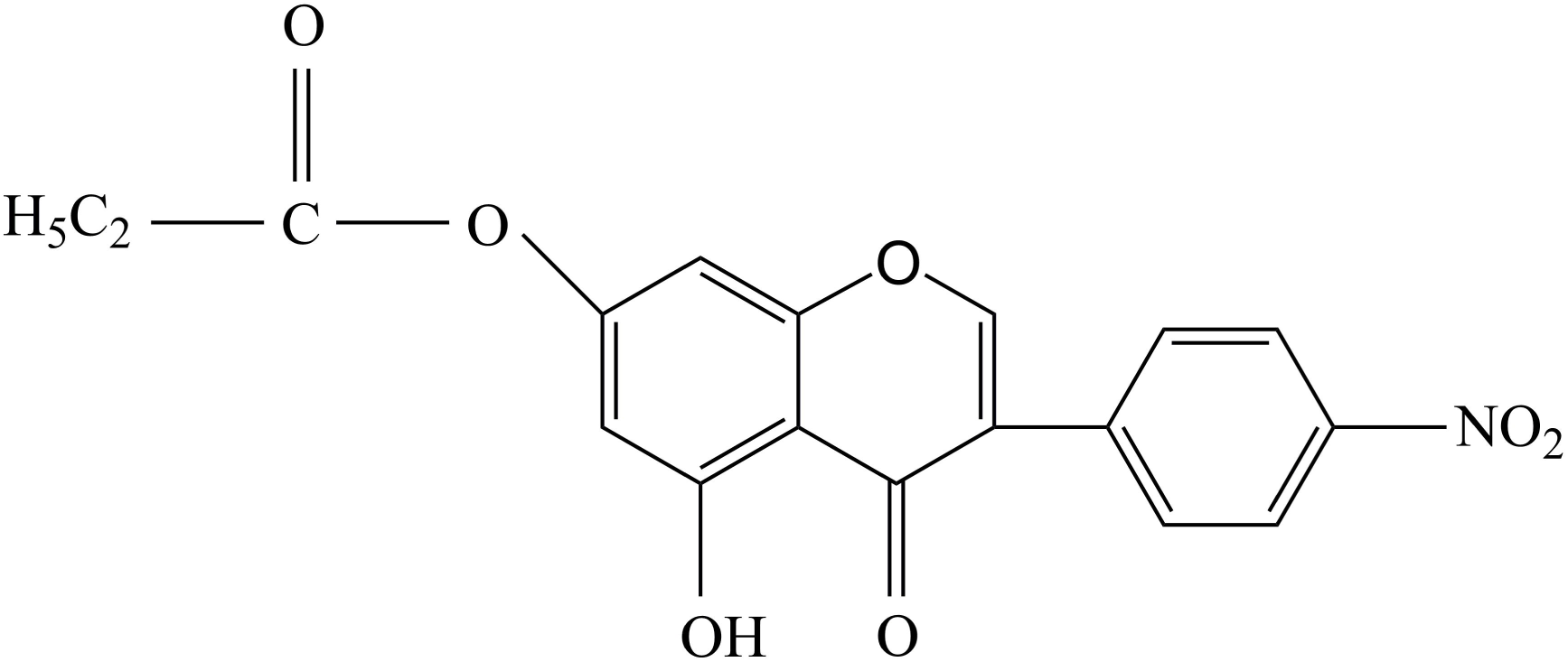
Chemical structure of 5-hydroxy-4’-nitro-7-propionyloxy-genistein.

**Figure 2.**
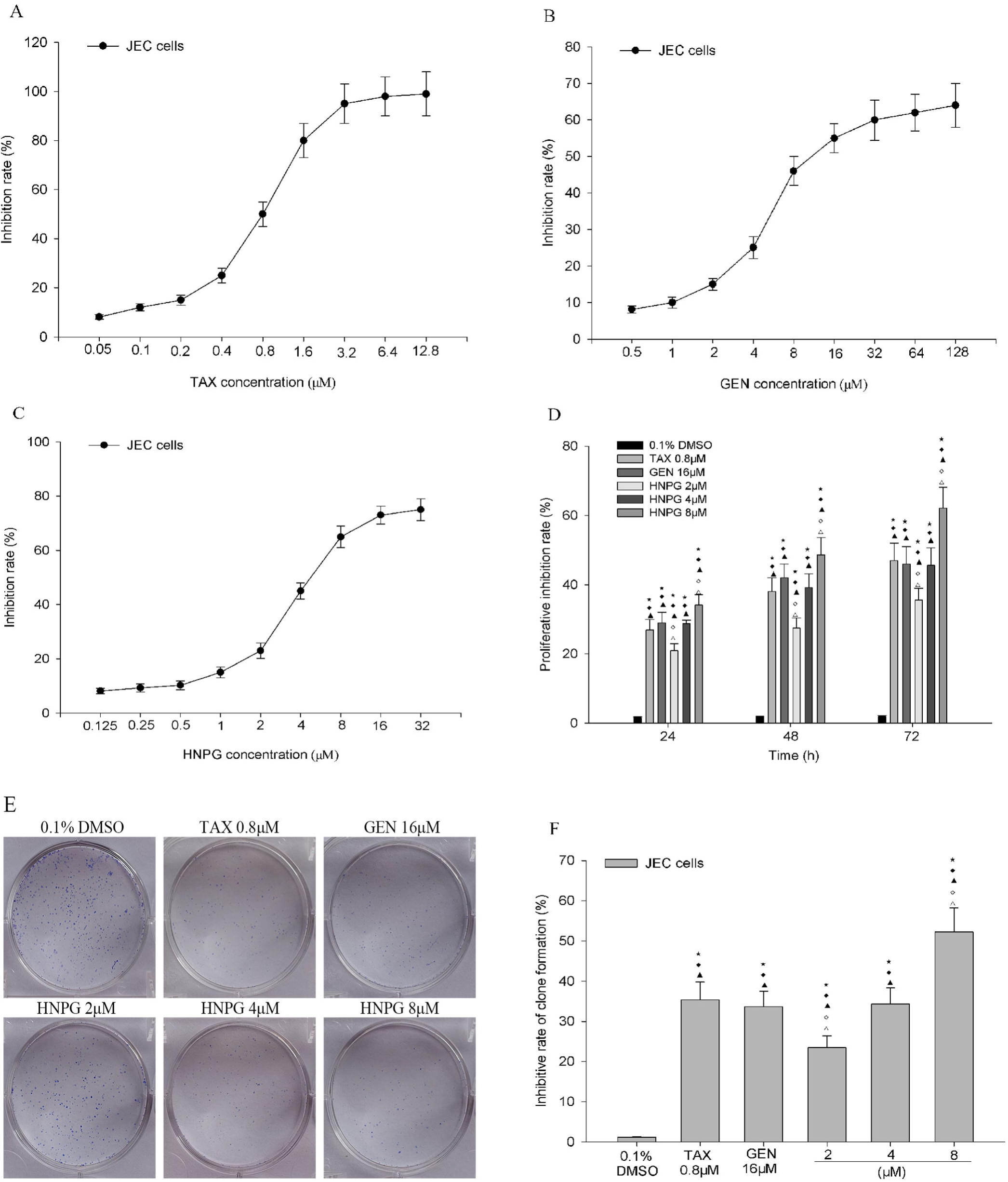
Effects of HNPG on the inhibition of cell viability in JEC cells. (A) Graph indicating the rate of proliferative inhibition of TAX ranging from 0.05 ~ 12.8 μM for 24 h. (B) Diagram showing the rate of proliferation inhibition of GEN ranging from 0.5 ~ 128 μM for 24 h. (C) Graph exhibiting the rate of proliferation inhibition of HNPG ranging from 0.125 ~ 32 μM for 24 h. (D) Histograms demonstrating the proliferative inhibition rate of 0.1 % DMSO, TAX of 0.8 μM, GEN of 16 μM and different concentrations of HNPG (2, 4 or 8 μM) for 24, 48 and 72 h. (E) Images indicating the clone formation of JEC cells exposed to 0.1 % DMSO, TAX of 0.8 μM, GEN of 16 μM and different concentrations of HNPG (2, 4 or 8 μM) for seven days stained with Giemsa stain. (F) Histogram demonstrating the inhibition rate of clone formation of JEC cells exposed to 0.1 % DMSO, TAX of 0.8 μM, GEN of 16 μM and different concentrations of HNPG (2, 4 or 8 μM) for seven days. The data are presented as the mean ± standard deviation from three independent experiments. ⋆P<0.05 vs. 0.1 % DMSO group, ♦P<0.05 vs. 0.8 μM TAX group,▲P<0.05 vs. 16 μM GEN group,♢P<0.05 vs. 2 μM HNPG group, ▵P<0.05 vs. 4 μM HNPG. HNPG, 5-hydroxy-4’-nitro-7-propionyloxy-genistein; TAX, taxol; GEN, genistein; DMSO, dimethyl sulfoxide.

### HNPG suppressed the clone formation of JEC cells

JEC cells were incubated at 37 °C with 0.1 % DMSO, TAX of 0.8 μM, GEN of 16 μM and different concentrations of HNPG (2, 4 or 8 μM) for seven days. The rate of clone formation was dramatically reduced and the cell amount inside the clones was significantly decreased. The inhibition rate of clone formation was significantly increased in a dose-dependent manner, every HNPG-treated group demonstrated a marked statistical difference compared with the control group (P_2μM/0.1%DMSO_<0.05, P_4μM/0.1%DMSO_<0.05, P_8μM/0.1%DMSO_<0.05). In addition, there was statistical difference among each HNPG-treated group (P_2/4μM_<0.05, P_2/8μM_<0.05, P_4/8μM_<0.05). While there was no statistical difference among TAX of 0.8 μM, GEN of 16 μM and HNPG of 4 μM (P_HNPG of 4μM/TAX of 0.8μM_>0.05, P_HNPG of 4μM/GEN of 16μM_>0.05, P_GEN of 16μM/TAX of 0.8μM_>0.05) (Fig. 2 E, F).

### HNPG inhibited the invasion ability of JEC cells

JEC cells were cultured with 0.1 % DMSO, TAX of 0.8 μM, GEN of 16 μM and different concentrations of HNPG (2, 4 or 8 μM) for 18 h, and the invasive ability was markedly decreased in a dose-dependent manner. The results demonstrated that the average cell numbers of 0.1 % DMSO, TAX of 0.8 μM, GEN of 16 μM and different concentrations of HNPG (2, 4 or 8 μM) groups invading through the Matrigel were 65.42 ± 5.64, 24.38 ± 3.16, 26.74 ± 3.26, 40.86 ± 4.71, 22.54 ± 3.26 and 12.37 ± 2.67, respectively. There was a statistical difference between each HNPG-treated group and the control (P_2μM/0.1%DMSO_<0.05, P_4μM/0.1%DMSO_<0.05, P_8μM/0.1%DMSO_<0.05), and there was statistical difference among each HNPG-treated group (P_2/4μM_<0.05, P_2/8μM_<0.05, P_4/8μM_<0.05), but there was no statistical difference among TAX of 0.8 μM, GEN of 16 μM and HNPG of 4 μM (P_HNPG of 4μM/TAX of 0.8μM_>0.05, P_HNPG of 4μM/GEN of 16μM_>0.05, P_GEN of 16μM/TAX of 0.8μM_>0.05) (Fig. 3 A, B).

**Figure 3.**
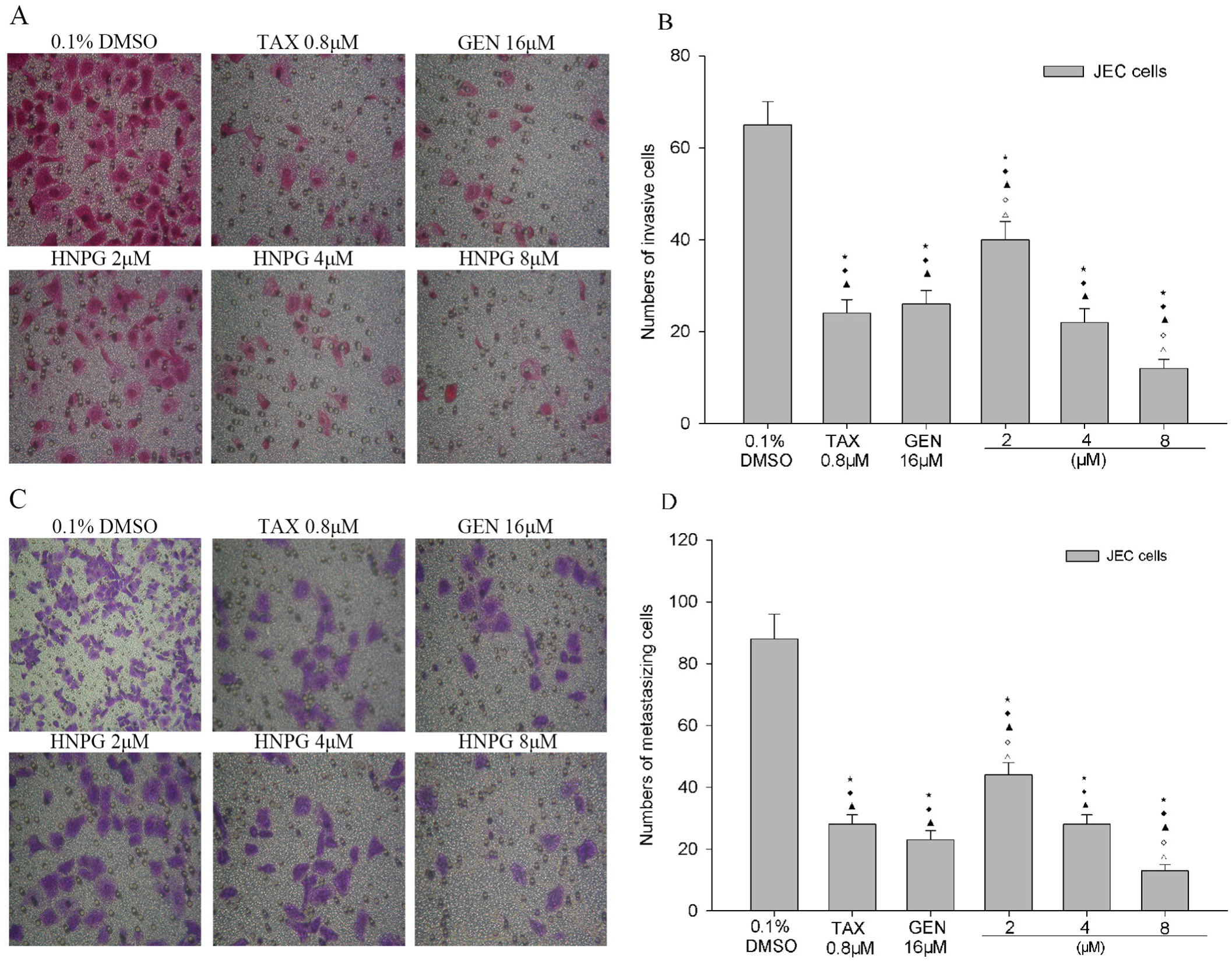
Effects of HNPG on the invasive and metastasizing capabilities of JEC cells treated with 0.1 % DMSO, TAX of 0.8 μM, GEN of 16 μM and different concentrations of HNPG (2, 4 or 8 μM) for 18 h. (A) Images demonstrating JEC cells in a Matrigel assay and stained with Hematoxylin and Eosin stain (magnification, 200 x). (B) Histogram exhibiting the numbers of invasive cells via a Matrigel assay. (C) Images indicating JEC cells on a polycarbonate membrane stained with crystal violet stain (magnification, 200 x). (D) Histogram showing the cell numbers of metastasis via a polycarbonate membrane. The data are presented as the mean ± standard deviation from three independent experiments. ϏP<0.05 vs. 0.1 % DMSO group, ♦ P<0.05 vs. 0.8 μM TAX group, ▴P<0.05 vs. 16 μM GEN group, ◊P<0.05 vs. 2 μM HNPG group, ▵P<0.05 vs. 4 μM HNPG. HNPG, 5-hydroxy-4’-nitro-7-propionyloxy-genistein; TAX, taxol; GEN, genistein; DMSO, dimethyl sulfoxide.

### HNPG suppressed the metastasis ability of JEC cells

JEC cells were cultivated with 0.1 % DMSO, TAX of 0.8 μM, GEN of 16 μM and different concentrations of HNPG (2, 4 or 8 μM) for 18 h, and the metastasizing ability of JEC cells was significantly decreased in a concentration-dependent manner. The results demonstrated that the average cell numbers of 0.1 % DMSO, TAX of 0.8 μM, GEN of 16 μM and different does of HNPG (2, 4 or 8 μM) groups that metastasized through polycarbonate membrane were 88.36 ± 7.42, 28.3.45 ± 3.82, 24.15 ± 3.21, 44.76 ± 4.18, 28.72 ± 3.18 and 13.24 ± 2.92, respectively. There was a statistical difference among every HNPG-treated group. There was a statistical difference between each HNPG-treated group and the control (P_2μM/0.1%DMSO_<0.05, P_4μM/0.1%DMSO_<0.05, P_8μM/0.1%DMSO_<0.05), and there was statistical difference among each HNPG-treated group (P_2/4μM_<0.05, P_2/8μM_<0.05, P_4/8μM_<0.05), while there was no statistical difference among TAX of 0.8 μM, GEN of 16 μM and HNPG of 4 μM (P_HNPG of 4μM/TAX of 0.8μM_>0.05, P_HNPG of 4μM/GEN of 16μM_>0.05, P_GEN of 16μM/TAX of 0.8μM_>0.05) (Fig. 3 C, D).

### HNPG induced the accumulation of G1 phase of JEC cells

JEC cells were exposed to 0.1 % DMSO, TAX of 0.8 μM, GEN of 16 μM, and different concentrations of HNPG (2, 4 or 8 μM) for 48 h, and G1 phase of JEC cells was significantly accumulated. The results indicated that HNPG could markedly block JEC cells in G1 phase in a dose-dependent manner, the G1 phase of 0.1 % DMSO, TAX of 0.8 μM, GEN of 16 μM, and different concentrations of HNPG (2, 4 or 8 μM) groups were 48.71 ± 2.12 %, 72.22 ± 4.15 %, 70.76 ± 5.52 %, 63.87 ± 4.56%, 72.05±5.21 % and 93.25 ± 6.54 %, respectively. Every HNPG-treated group possessed statistical difference compared with control group (P_2μM/0.1%DMSO_<0.05, P_4μM/0.1%DMSO_<0.05, P_8μM/0.1%DMSO_<0.05), and there were significant differences among each HNPG-treated group (P_2/4μM_<0.05, P_2/8μM_<0.05, P_4/8μM_<0.05), there were no difference among TAX of 0.8 μM, GEN of 16 μM and HNPG of 4 μM (P_TAX of 0.8μM/HNPG of 4μM_>0.05, P_GEN of 16μM/HNPG of 4μM_>0.05, P_GEN of 16μM/TAX of 0.8μM_>0.05) (Fig. 4 A, B).

**Figure 4.**
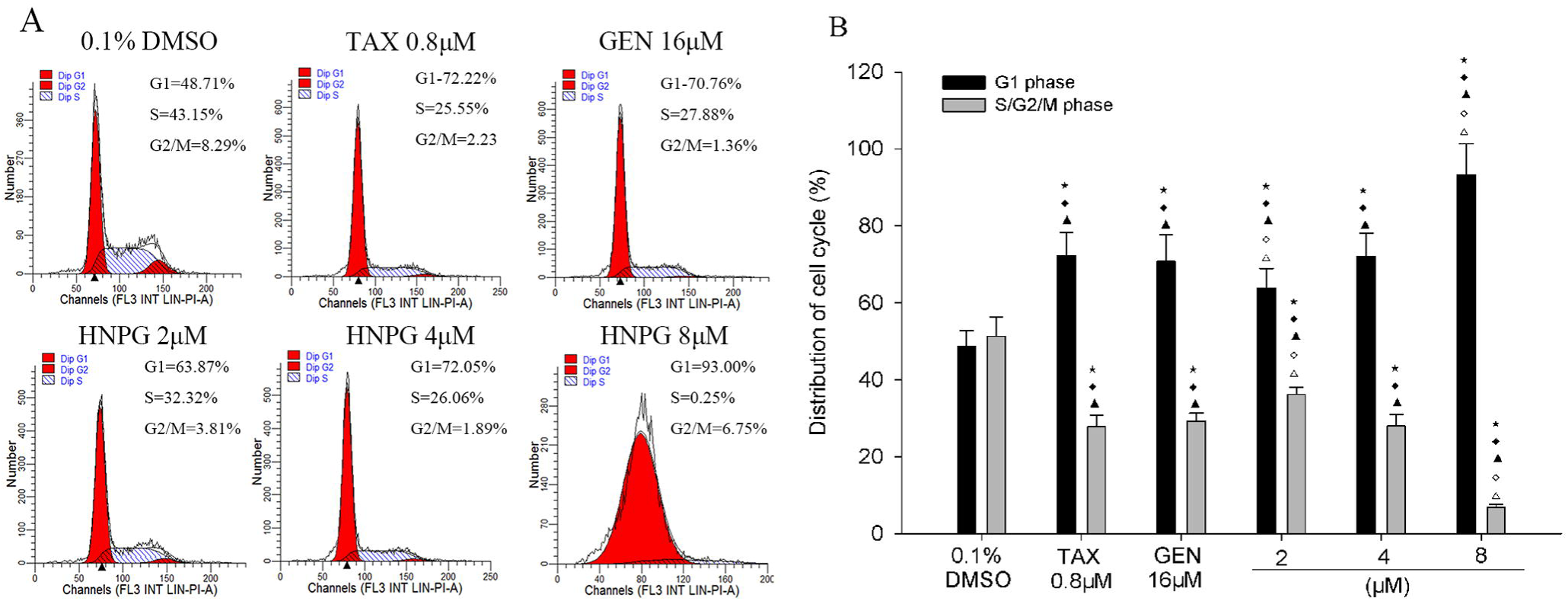
Effects of HNPG on the accumulation in G1 phase in JEC cells cultured with 0.1 % DMSO, TAX of 0.8 μM, GEN of 16 μM and different concentrations of HNPG (2, 4 or 8 μM) for 48 h. (A) Diagrams exhibiting the distribution of cell cycle percentage of JEC cells stained with PI. (B) Histogram demonstrating the distribution of cell cycle percentage rate. The data are presented as the mean ± standard deviation from three independent experiments. ⋆P<0.05 vs. 0.1 % DMSO group, ♦P<0.05 vs. 0.8 μM TAX group, ▴P<0.05 vs. 16 μM GEN group, ♢P<0.05 vs. 2 μM HNPG group, ▵P<0.05 vs. 4 μM HNPG. HNPG, 5-hydroxy-4’-nitro-7-propionyloxy-genistein; TAX, taxol; GEN, genistein; DMSO, dimethyl sulfoxide; PI, propidium iodide.

### HNPG changed the morphological features of JEC cells

The morphology of cells of 0.1 % DMSO group was long fusiform, the filamentous pseudopod was distributed around the cell body, and the surface of the cell membrane was relatively flat covered with granules in 60 μm^2^ × 60 μm^2^ of scan range of AFM (Fig. 5 A, B). The surface of the cell membrane showed the shape of flannelette blanket distributing small homogeneous bulges, the arrangement of bulges was moderate in 5 μm^2^ × 5 μm^2^ of scanning range, which is characterized by normal malignant tumor cells (Fig. 5 C, D). When cells were treated by 0.1 % DMSO, TAX of 0.8 μM, GEN of 16 μM and different concentrations of HNPG (2, 4 or 8 μM) after 24 h, the morphology of JEC cells obviously changed in concentration-dependent manner, the cell body retracted, the filamentary pseudopod disappeared, the surface of the cell membrane appeared to holes of unequal size, the nucleus collapsed, the height of nucleus decreased, and ultrastructure showed there was irregular bumps around the hole of the cell membrane which indicated that the cells were of obvious damage. The degree of damage caused by HNPG of 40 μM was approximately similar to TAX of 0.8 μM or GEN of 16 μM (Fig. 5 A, B, C, D).

**Figure 5.**
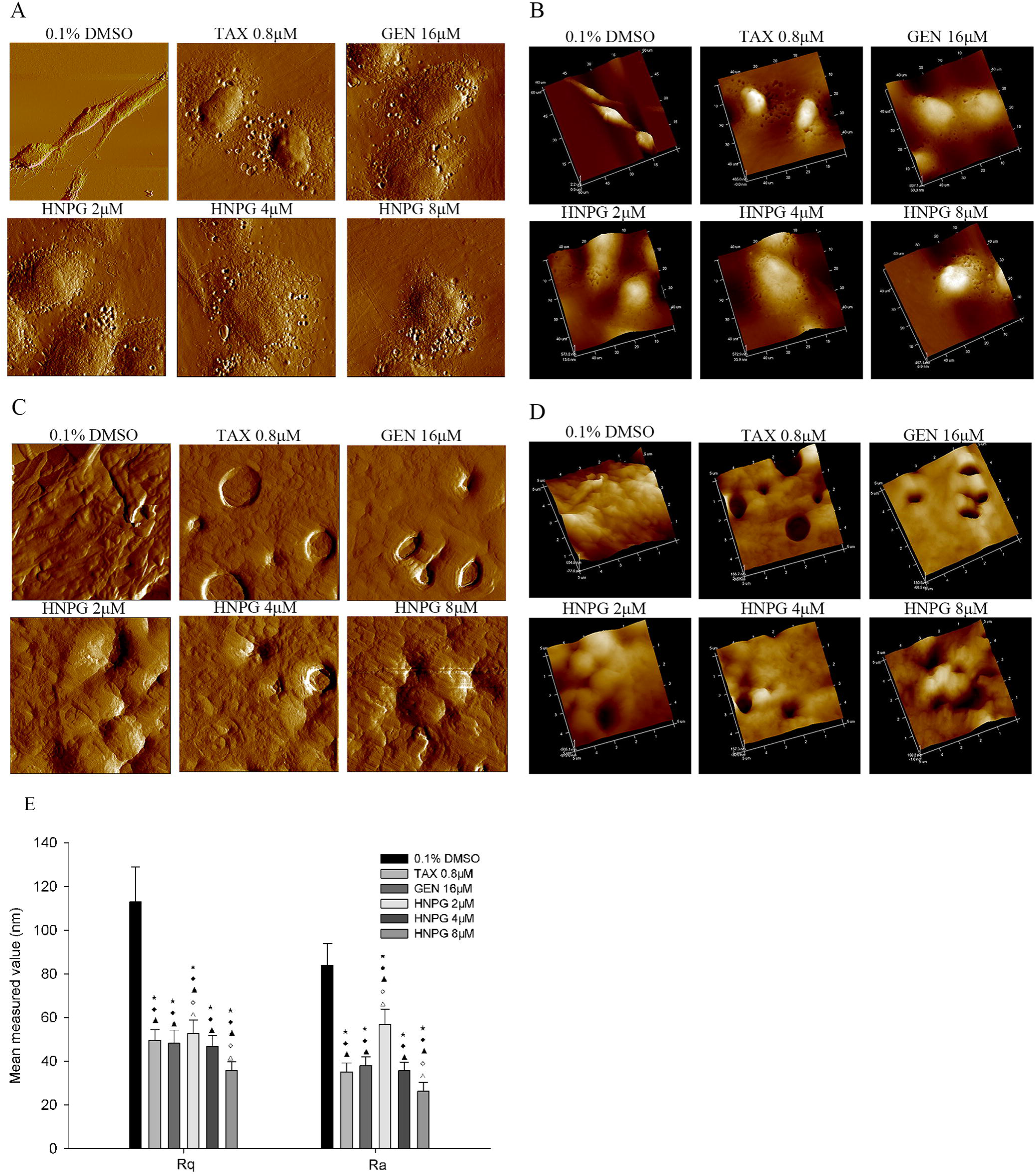
The changes of morphological features and the detection of biological force trait using atomic mechanics microscopy (AFM) in JEC cells incubated with 0.1 % DMSO, TAX of 0.8 μM, GEN of 16 μM and different concentrations of HNPG (2, 4 or 8 M) for 24 h. (A) Images exhibiting the morphological characteristics of two-dimensional diagram of JEC cells in 60 μM^2^ × 60 μM^2^ of scan range. (B) Images demonstrating the morphological features of three-dimensional diagram of JEC cells in 60 μM^2^ × 60 μM^2^ of scan range. (C) Images indicating the morphological characteristics of two-dimensional diagram of JEC cells in 5 m^2^ × 5 m^2^ of scan range. (D) Images showing the morphological features of three-dimensional diagram of JEC cells in 5 μM^2^ × 5 μM^2^ of scan range. (E) Histogram exhibiting the changes of root-mean-square roughness (Rq) and average roughness (Ra) of JEC cells. (F) Histograms demonstrating the changes of Young's Modulus of Gaussian fitting of JEC cells. (G) Histograms indicated the changes of the adhesive force of Gaussian fitting of JEC cells. The data are indicated as the mean ± standard deviation from three independent experiments. ⋆P<0.05 vs. 0.1 % DMSO group, ♦P<0.05 vs. 0.8 μM TAX group, ▴P<0.05 vs. 16 μM GEN group, ◊P<0.05 vs. 2 μM HNPG group, ▵P<0.05 vs. 4 μM HNPG. HNPG, 5-hydroxy-4’-nitro-7-propionyloxy-genistein; TAX, taxol; GEN, genistein; DMSO, dimethyl sulfoxide.

### HNPG decreased the root-mean-square roughness (Rq) and the average roughness (Ra) of JEC cells

The present study found that the root-mean-square roughness (Rq) of 0.1 % DMSO, TAX of 0.8 μM, GEN of 16 μM and different concentrations of HNPG (2, 4 or 8 μM) was 113.0 ± 16.11 nm, 49.557 ± 5.40 nm, 48.33 ± 5.66 nm, 52.90 ± 5.80 nm, 46.87 ± 6.64 nm and 35.77 ± 4.24 nm respectively. In addition, the average roughness (Ra) of 0.1 % DMSO, TAX of 0.8 μM, GEN of 16 μM and different concentrations of HNPG (2, 4 or 8 μM) was 83.97 ± 7.82 nm, 35.2 ± 4.5 nm, 37.97 ± 5.13 nm, 56.93 ± 6.48 nm, 35.73 ± 4.72 nm and 26.43 ± 2.33 nm respectively. The Rq and Ra of every HNPG-treated group possessed statistical difference compared with control group (P_2μM/0.1%DMSO_<0.05, P_4μM/0.1%DMSO_<0.05, P_8μM/0.1%DMSO_<0.05), and there were significant differences among the Rq and Ra of each HNPG-treated group (P_2/4μM_<0.05, P_2/8μM_<0.05, P_4/8μM_<0.05), there were no difference among the Rq and Ra of TAX of 0.8 μM, GEN of 16 μM and HNPG 4 μM (P_TAX of 0.8μM/HNPG of 4μM_>0.05, P_GEN of 16μM/HNPG of 4μM_>0.05, P_GEN of 16μM/TAX of 0.8μM_>0.05) (Fig. 5 E).

### HNPG increased Young's modulus and adhesion force of Gaussian fitting of JEC cells

The mechanical detection of individual living cell was measured using AFM indentation technique, and the value of Young's modulus of Gaussian fitting was 7.53 ± 1.12 kPa, 17.72 ± 2.55 kPa, 18.39 ± 2.97 kPa, 12.53 ± 1.14 kPa, 18.18±2.73 kPa and 20.46 ± 2.15 kPa respectively. The nonspecific forces between the cell surface and the probe was detected using AFM indentation technique, and the adhesion force of Gaussian fitting between the cell surface and the probe was 53.23 ± 4.01 pN, 69.36 ± 5.36 pN, 69.10 ± 5.03 pN, 62.62 ± 7.97 pN, 73.49 ± 5.23 pN and 80.19 ± 12.10 pN respectively. Young's modulus and adhesion force of Gaussian fitting of every HNPG-treated group possessed statistical difference compared with control group (P_2μM/0.1%DMSO_<0.05, P_4μM/0.1%DMSO_<0.05, P_8μM/0.1% DMSO_<0.05), and there were significant differences among Young's modulus and adhesion force of Gaussian fitting of each HNPG-treated group (P_2/4μM_<0.05, P_2/8μM_<0.05, P_4/8μM_<0.05), there were no difference among Young's modulus and adhesion force of Gaussian fitting of TAX of 0.8 μM, GEN of 16 μM and HNPG 4 μM (P_TAX of 0.8μM/HNPG of 4μM_>0.05, P_GEN of 16μM/HNPG of 4μM_ <0.05, P_GEN of 16μM/TAX of 0.8μM_<0.05) (Fig. 6 A, B).

**Figure 6.**
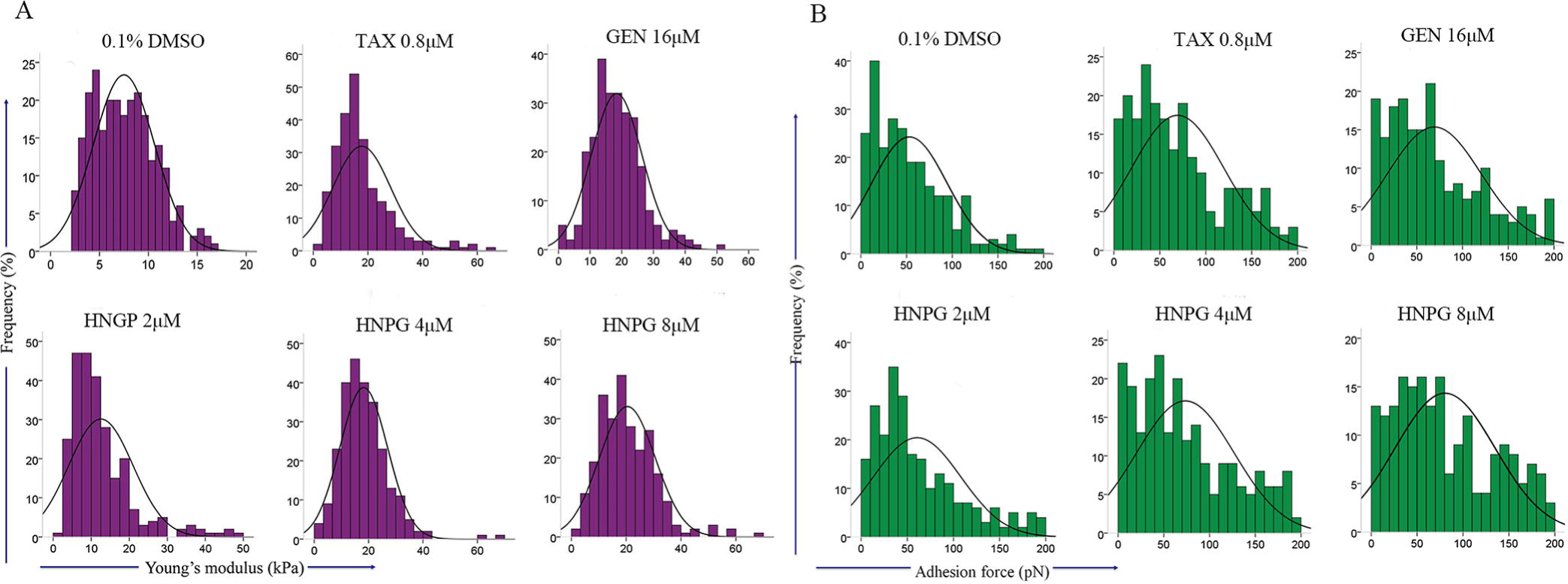
The biological force trait using atomic mechanics microscopy (AFM) in JEC cells incubated with 0.1 % DMSO, TAX of 0.8 μM, GEN of 16 μM and different concentrations of HNPG (2, 4 or 8 μM) for 24 h. (A) Histograms demonstrating the changes of Young's Modulus of Gaussian fitting of JEC cells. (B) Histograms indicated the changes of the adhesive force of Gaussian fitting of JEC cells. The data are indicated as the mean ± standard deviation from three independent experiments.

### HNPG regulated mRNA of wnt/β-catenin signal pathway in JEC cells

JEC cells were administrated to 0.1 % DMSO and different concentrations of HNPG (2, 4 or 8 μM) for 24 h, and the average value of mRNA of β-catenin, C-myc, Cyclin D1, MMP-2, MMP-7 and MMP-9 were analyzed. The expressive mRNA level of β-catenin, C-myc, Cyclin D1, MMP-2, MMP-7 and MMP-9 exhibited a decreasing trend in dose-dependent, every HNPG-treated groups exhibited a statistical difference compared with the control group (P_2μM/0.1%DMSO_<0.05, P_4μM/0.1%DMSO_<0.05, P_8μM/0.1%DMSO_<0.05) in β-catenin, C-myc, Cyclin D1, MMP-2, MMP-7 and MMP-9 mRNA expression levels. There were statistical differences among every HNPG-treated group in β-catenin, C-myc, Cyclin D1, MMP-2, and MM-7 and MMP-9 mRNA expression levels (P_2/4μM_<0.05, P_2/8μM_<0.05, P_4/8μM_<0.05) (Fig. 7 A, B).

**Figure 7.**
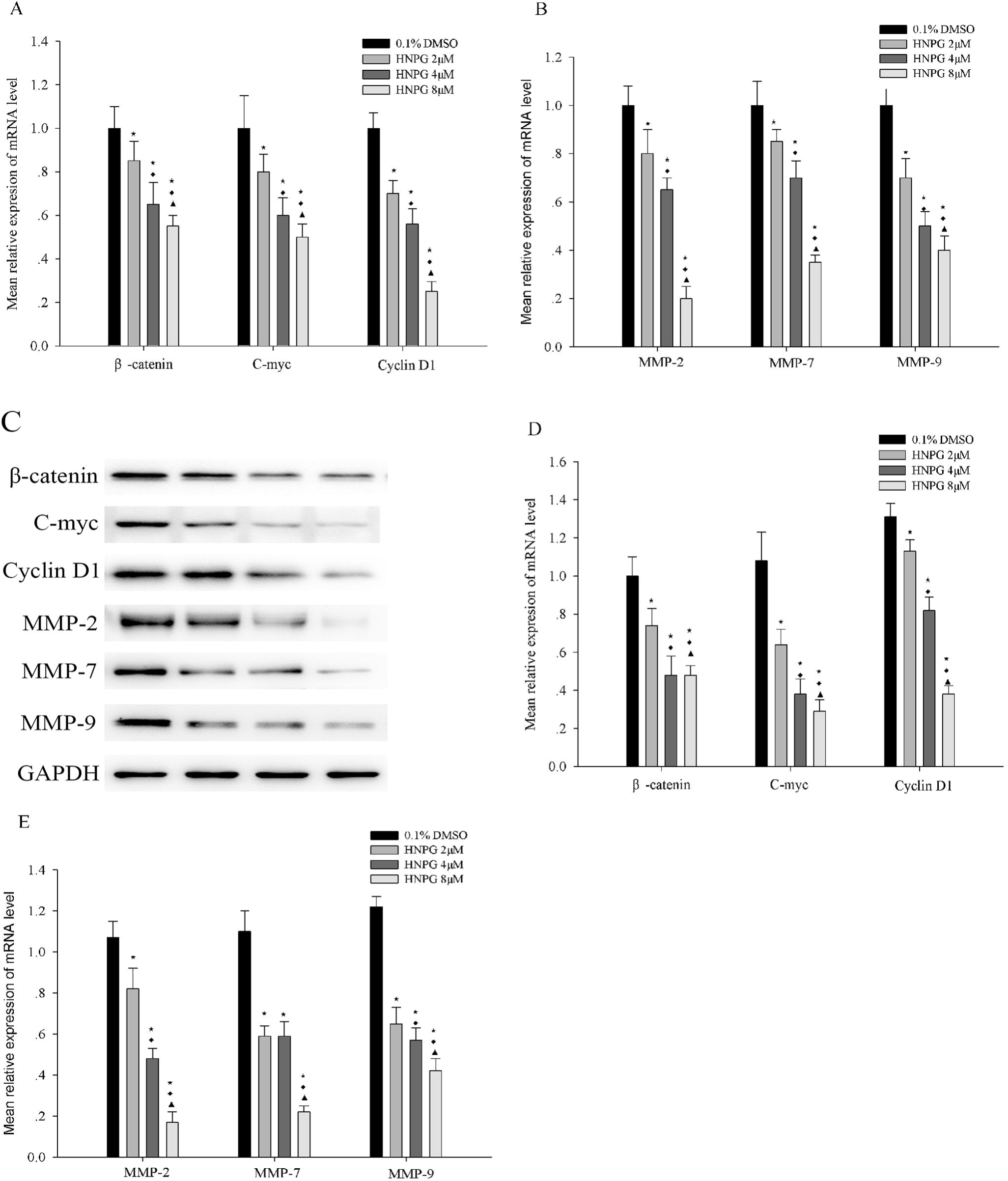
The changes of gene expression of JEC cells incubated with 0.1 % DMSO, and different concentrations of HNPG (2, 4 or 8 μM) for 24 h. (A) Histogram demonstrating the average relative expression of mRNA level of β-catenin, C-myc and Cyclin D1. (B) Histogram indicating the mean relative expression of mRNA of MMP-2, MMP-7 and MMP-9. (C) Electrophoretograms showing the expression of protein level of β-catenin, C-myc, Cyclin D1, MMP-2, MMP-7 and MMP-9. (D) Histogram demonstrating the average relative expression of protein level of β-catenin, C-myc and Cyclin D1. (E) Histogram exhibiting the mean relative expression of protein level of MMP-2, MMP-7 and MMP-9. The data were shown as the mean ± standard deviation from three independent experiments. ⋆P<0.05 vs. 0.1 % DMSO group, ♦ P<0.05 vs. 2 μM HNPG group, ▲P<0.05 vs. 4 μM HNPG. HNPG, 5-hydroxy-4’-nitro-7-propionyloxy-genistein; DMSO, dimethyl sulfoxide.

### HNPG regulated protein of wnt/β-catenin signal pathway of JEC cells

JEC cells were exposed to 0.1 % DMSO and different concentrations of HNPG (2, 4 or 8 μM) for 24 h, and the average relative density of β-catenin, C-myc, Cyclin D1, MMP-2, MMP-7 and MMP-9 were analyzed. The expression level of β-catenin, C-myc, Cyclin D1, MMP-2, MMP-7 and MMP-9 protein exhibited a decreasing trend in concentration-dependent. Every HNPG-treated groups demonstrated a statistical difference compared with the control group (P_2μM/0.1%DMSO_<0.05, P_4μM/0.1%DMSO_<0.05, P_8μM/0.1%DMSO_<0.05) in β-catenin, C-myc, Cyclin D1, MMP-2, MMP-7 and MMP-9 protein expression levels. There were statistical differences among every HNPG-treated group in β-catenin, C-myc, Cyclin D1, MMP-2 and MMP-9 protein expression levels (P_2/4μM<0.05_, P_2/8μ_M<0.05, P_4/8μM_<0.05). There were significant differences between 2, or 4 μM HNPG-treated group and 8 μM HNPG-treated groups in MMP-7 protein expression (P_2/8μM_<0.05 or P_4/8μM_<0.05), but there were no difference between 2 and 4 μM HNPG-treated groups in MMP-7 protein expression (P_2/4μM_>0.05) (Fig. 7 C, D, E).

## DISCUSSION

Invasion and metastasis are basic characteristics of malignant tumors, malignant carcinomas not only germinate in the primary site via infiltrating and damaging adjacent organs and tissues, but also metastasize to other areas to proliferate and grow and cause new damage, so the capacities of invasion and metastasis directly reflect the malignant degree of tumors, and the capacity of inhibiting invasion and metastasis are also becoming one of the normal indices to evaluate the pharmacological activity (Shapiro et al., 2017; Chen et al., 2017). Previous studies have demonstrated that HNPG could inhibit proliferation in gastric and breast cancer cells *in vitro* (Wang et al., 2012; Wang et al., 2005), but the therapeutic effect of HNPG has not been compared with its precursor GEN and clinical chemotherapy drugs, and its molecular mechanism of inhibition of proliferation has not yet been elucidated. In the present study, the data indicated that HNPG could suppress proliferation, clone formation, invasion and metastasis, and accumulated G1 phase in human endometrial cancer JEC cells *in vitro* compared with GEN and TAX. These experimental results suggested HNPG may be an excellent novel candidate for therapy in human endometrial cancer.

Cells proliferate via cell cycle, one cell dividing into two cells has to go through first gap phase (G1), synthetic phase (S), second gap phase (G2) and mitotic phase (M) in sequence, and any cell phases encounter obstacles will result in cell proliferation stasis even apoptosis or necrosis, which will affect the function of cell viability such as proliferation, colony formation, invasion and metastasis (Kim et al., 2017). When external factors and internal factors cause cell cycle damage, the permeability of cell membrane will be increased, which made propidium iodide (PI) enter the cytoplasm through the cell membrane, and combine with DNA in the nucleus (So et al., 2017). The content of DNA in different cell phases is different, the feature of DNA in G1 phase is characterized in diploid cells, the trait of DNA in G2 phase is represented in tetraploid cells, while the characteristics of DNA in S phase is between diploid cells and tetraploid cells. Different cell phases will be distinguished via detecting the intensity of PI fluorescence combing with DNA by flow cytometry (Kim et al., 2017; So et al., 2017). In the results of the present study, the numbers of G1 phase of JEC cells were markedly enhanced in a dose-dependent manner, following HNPG treatment for 48 h. Therefore, it was suggested that the HNPG-mediated inhibition of proliferation, clone formation, invasion and metastasis of JEC cells might occur via the accumulation in G1 phase.

It is well-known cell proliferation has to go through check points which are regulated by cyclin, cyclin dependent kinase (CDK) and cyclin dependent kinase inhibitor (CKI) (Kim et al., 2018). The check point of G1/S is one of most important check points in the progress of cell cycle, when the check point of G1/S receive the signal information of DNA damage, it will interrupt the progress of cell cycle and block cell in G1 phase (Kim et al., 2018). Cyclin D1 is most important regulatory factor in G1/S, and it can combine with CDK2, CDK4 and CDK6, and promote cells go through G1/S check point and enter S phase (Kim et al., 2018). Once the amount of Cyclin D1 increase in cells, which will accelerate the proliferative speed and cause tumor and other diseases, conversely if the amount of Cyclin D1 decreases, cells will accumulate in G1 phase, which will result in the cell function decline, including invasion and metastasis (Kim et al., 2018; Morita et al., 2017). C-myc can promote cell proliferation via regulating several gene related to proliferation including Cyclin D1 and so on (Morita et al., 2017). It is reported that C-myc and Cyclin D1 are closely related to tumorigenesis, invasion and metastasis (Kim et al., 2018; Morita et al., 2017). In the present study, C-myc and Cyclin D1 of JEC cells incubated with different concentrations of HNPG were notably down-regulated in dose-dependent manner, accompanied by the accumulation of G1 phase, which suggested the accumulation of G1 phase might be related to C-myc and Cyclin D1 down-regulated.

Degradation of extracellular matrix (ECM) is an important element of invasion and metastasis of malignant tumors, and matrix metalloproteinases (MMPs) is a group of proteolytic enzymes, which is closely related to various pathological processes, especially tumor’s invasion and metastasis (Rossi et al., 2017; Taniguchi et al., 2017). MMP-7, MMP-2 and MMP-9 are important members of the family of MMPs, can degrade extracellular matrix and basement membrane components, and it is reported MMP-7, MMP-2 and MMP-9 are up-regulated in tumor tissues, regulating tumor cell proliferation and metastasis through degradation of ECM (Rossi et al., 2017; Taniguchi et al., 2017). MMP-2 and MMP-9 are IV type collagen enzyme, mainly degrade IV type collagen and laminin (Rossi et al., 2017). MMP-7 is a special type MMP with strong degrading activity of ECM and possessing wide range of biological substrates including type collagen, laminin, polysaccharide, type gelatin, soluble elastin and so on (Taniguchi et al., 2017). In our research, we found HNPG inhibited invasion and metastasis of JEC cells accompanied with MMP-2, MMP-7 and MMP-9 down-regulated in dose-dependent manner, which suggested that MMP-2, MMP-7 and MMP-9 were involved in the effects of HNPG on inhibiting invasion and metastasis.

Adhesion force decreased among cells and amoeba movement depending on pseudopod are the premise conditions of invasion and metastasis in malignant cancer cells (Cascione et al., 2017). Cells may secrete certain adhesive proteins that fasten cells together and prevent cells from metastasizing to other sites (Cascione et al., 2017). The force produced by those adhesive proteins can be measured using AFM, the adhesive force of malignant cancer tissue is lower than normal tissue, so measuring the size of adhesive force between cells can reflect the metastasizing capacities of tumor cells (Meng et al., 2017). The plasticity or elasticity of cells is the necessary condition of cells migration, Young's modulus is often used to detect the plasticity or elasticity of cells in bioengineering (Laurito et al., 2016). In the present study, we found JEC cells treated by different concentration of HNPG became oval, pseudopod disappeared, the surface roughness (Rq and Ra) decreased, while adhesion force and Young's modulus of Gaussian fitting increased, which were in conjunction with the inhibition of invasion and metastasis. Our researches suggested the inhibition effects of HNPG on invasion and metastasis was related to the alteration of morphology, the decrease of Rq and Ra, and the increase of Young's modulus and adhesive force.

Wnt/β-catenin signal pathway is involved in many carcinogenesis, invasion and metastasis, and plays a very important role in tumor chemotherapy including endometrial cancer (Laurito et al., 2017). β-catenin occupy the core position in wnt/β-catenin signal pathway, once the amount of β-catenin increased in cytoplasm, β-catenin will enter nucleus, which will trigger a series of biological reactions, promote the expression of C-myc, Cyclin D1, and MMP-7, MMP-2 and MMP-9, lead to a lot of cellular events including proliferation, invasion and metastasis and so on (Kasprzak et al., 2017). In the present study, we found HNPG caused the inhibition of proliferation, invasion, metastasis, accompanied by the inactivation of β-catenin, which suggested that wnt/β-catenin signal pathway was involved in the pharmacological effects of HNPG on inhibiting cell viability of JEC cells.

In conclusion, HNPG demonstrated significant cytotoxic activity in human endometrial carcinoma JEC cells. HNPG inhibited proliferation, clone formation, invasion and metastasis, and accumulated cell cycle in G1 phase *in vitro*. Simultaneously, Rq and Ra decreased, Young's modulus and adhesion force increased in JEC cells. Additionally, HNPG inactivated wnt/β-catenin signal pathway, down-regulated the expression of β-catenin, C-myc, Cyclin D1, MMP-2, MMP-7 and MMP-9, the present study not only detected the inhibiting effects of cell viability, but also elucidated the possible molecular mechanism of HNPG. In summary, HNPG indicated a marked cytotoxic activity in human endometrial carcinoma JEC cells via inactivating wnt/β-catenin signal pathway, which suggested that HNPG may be a novel candidate for chemotherapeutic drug. Subsequent studies will focus on investigating the metabolism of HNPG in experimental animal models, detecting its blood drug concentration and its half-life and the side effects on the brain, heart, lung, liver and kidney cells, and elucidate the possible molecular mechanism *in vivo*.

## MATERIALS AND METHODS

### Materials

5-hydroxy-4’-nitro-7-propionyloxy-genistein (HNPG) and genistein (GEN) were purchased from Department of Organic Chemistry, School of Pharmacy, Second Military Medical University (Shanghai China). HNPG is a kind of pale yellow crystalline powder, and its molecular formula is C_18_H_13_O_7_N with a molecular weight of 355, and its chemical structure is presented in Fig 1. HNPG and GEN were dissolved in dimethyl sulfoxide (DMSO) into 0.1 % (v/v) stock solution until use, which concentration was made sure there was no effect on cell proliferation. Taxol (TAX) was bought from Molbase technology Co., Ltd. (Catalog Number: SJ0033A332183; Wuhan, Hubei, China). DMSO, PBS and MTT obtained from Tiangen Biotech Co., Ltd. (Beijing, China). Giemsa, crystal violet and hematoxylin & eosin (H&E) stains were all bought form Sangon Biotech Co., Ltd. (Shanghai, China). Matrigel™ was purchased from Shanghai Invitrogen Biological Technology Co., Ltd. (Shanghai, China). Specific antibodies for rabbit anti-β-catenin (Catalog Number: A00183), rabbit anti-C-myc (Catalog Number: D110006) and rabbit Cyclin D1 (Catalog Number: D151941) obtained from Sangon Biotech Co., Ltd. (Shanghai, China). Rabbit anti-MMP-2 (Catalog Number: M00286), rabbit anti-MMP-7 (Catalog Number: PB0071), rabbit anti-MMP-9 (Catalog Number: BA2202), rabbit anti-GAPDH antibodies (Catalog Number: A00227-1) and horseradish peroxidase-conjugated secondary rabbit antibody (Catalog Number: BA1082) were obtained from Boster Biotech Co., Ltd. (Wuhan, Hubei, China).

### Cell culture

JEC cells were purchased from the cell library of the Chinese Academy of Science (Shanghai, China) and cultured in RPMI-1640 (Thermo Fisher Scientific Inc., Waltham, MA, USA) that contained 10 % fetal bovine serum (FBS; HyClone™, GE Healthcare, Logan, UT, USA) and antibiotic antimycotic solution (100 x; Mediatech, Inc., Manassas, VA, USA) at 37 °C in 5 % CO_2_ humidified incubator. The cells were then divided in six groups: Control group (0.1 % DMSO), TAX group, GEN group and different concentrations of HNPG (2, 4 and 8 μM) groups.

### MTT assay

JEC cells were seeded into a 96-well plate (Tiangen Biotech Co., Ltd., Beijing, China) at a density of 1 × 10^4^ cells/well and incubated with different concentrations of TAX of 0.05, 0.1, 0.2, 0.4, 0.8, 1.6, 3.2, 6.4 and 12.8 μM, GEN of 0.5, 1, 2, 4, 8, 16, 32, 64 and 128 μM, and HNPG of 0.125, 0.25, 0.5, 1, 2, 4, 8, 16 and 32 μM for 24 h at 37 °C in 5 % Co_2_. Subsequently, 20 μl 5 mg/ml MTT stock solution was added to each well and continued to culture for 6 h at 37 °C. A total of 100 μl DMSO was used to stop the reaction, and spectrophotometric absorbance was subsequently measured using a microplate reader (ELX-800; Sangon Biotech Co., Ltd., Shanghai, China) at 570 nm (A570). The values of half maximal inhibitory concentration (IC50) of TAX, GEN and HNPG was obtained according to the value of spectrophotometric absorbance. TAX of 0.8 μM, GEN of 16 μM and HNPG (2, 4 and 8 IM) were used to the subsequent experiments to measure the inhibitive rates for 24, 48 and 72 h. The inhibitive rate (IR) was calculated as follows: (1-average A570 of the experimental group/average A570 of the control group) × 100 %. Experiments were performed in triplicate, and the mean value was calculated.

### Flat plate clone formation method

JEC cells were collected and seeded into a 6-well plate (Tiangen Biotech Co., Ltd., Beijing, China) at a density of 2 × 10^3^ cells/well and incubated for 24 h at 37 °C in 5 % CO_2_. Then 0.1 % DMSO, TAX of 0.8 μM, GEN of 16 μM and different concentrations of HNPG (2, 4 or 8 μM) were added to each well, and cultured for seven days at 37 °C in 5 % CO_2_ until visible clones formed. Clones containing >50 cells were defined as one clone, and were fixed with 95 % methanol for 15 min at 20 °C and stained with 0.1 % Giemsa stain for 15 min at 20 °C. The numbers of colon were computed and the clone formation inhibition rate (%) was calculated as follows: 1 - (the mean number of HNPG group/the mean number of control group) × 100 %. The results were representative of three independent experiments.

### Transwell assay

A pre-cooled 24-well Transwell plate (Tiangen Biotech Co., Ltd., Beijing, China) was covered with 30 μl Matrigel™ at a 1:3 dilution and incubated at 37 °C for 3 h. Then, JEC cells (1 × 10^5^/well) were cultivated at 37 °C for 18 h in the inner chamber and exposed to 100 μl 0.1 % FBS/RPMI-1640 with 0.1 % DMSO, TAX of 0.8 μM, GEN of 16 μM and different concentrations of HNPG (2, 4 or 8 μM), and the outer chamber was filled with 500 μl 10 % FBS/HNPG-1640 to act as a chemoattractant. Subsequently the invasive cells were fixed with 95 % ethanol for 15 min at 20 °C, and then stained with 0.5 % H&E stain for 15 min at 20 °C. JEC cells (1 × 10^5^/well) were seeded into the inner chamber of 24-well Transwell plate, treated with 100 μl 0.1 % FBS/RPMI-1640 containing 0.1 % DMSO, TAX of 0.8 μM, GEN of 16 μM and different concentrations of HNPG (2, 4 or 8 μM), while the outer chamber contained 500 μl 10 % FBS/RPMI-1640, and the Tanswell instrument were cultured for 18 h at 37 °C. Following this, the metastasized cells were fixed with 4 % paraformaldehyde for 15 min at 20 °C and subsequently stained with 0.1 % crystal violet stain for 15 min at 20 °C. The number of invasive cells or metastasized cells was manually counted in five randomly selected fields under inverted microscope (Type: N-STORM 4.0; Nikon instruments co. Ltd., Shanghai, China) with magnification 200 x. Every group had three repeat wells and the average value was calculated.

### Cell cycle analysis

JEC cells were treated with 0.1 % DMSO, TAX of 0.8 μM, GEN of 16 μM and different concentrations of HNPG (2, 4 or 8 μM) for 48 h, then rinsed with PBS twice, digested with 0.25 % trypsin, centrifuged 800 rpm for 5 min at 20 °C, discarded supernate, fixed in 70 % ethanol at 4 °C. Subsequently, the cells were stained with PI (50 mg/ml) solution (Tiangen Biotech Co., Ltd., Beijing, China) at 20 °C for 15 min in the darkness, and then analyzed using flow cytometry for distribution of cell cycle phase. Excitation and emission wavelengths of 488 nm were selected, experiments were performed in triplicate, and the mean value was calculated.

### Atomic force microscope (AFM) measurement

1 × 10^8^ JEC cells in logarithmic phase were seeded into 35 mm cell culture dish, and cultured in 0.1 % DMSO, TAX of 0.8 μM, GEN of 16 μM and different concentrations of HNPG (2, 4 or 8 μM) for 24 h, then fixed cells for 6 min at 20 °C using 4 % paraformaldehyde, and rinsed two times with PBS buffer, then the changes of morphology were observed and imaged and the Rq and Ra were detected using atomic force microscope (NanoScope IIIa MultiMode; Veeco Inc., New York, US). The radius of microcantilever probe was less than 10 nm, the force elastic coefficient of microcantilever was 5 N/m, the length or width of microcantilever was 125 μm or 25 μm respectively. The scan ranged from 60 μm^2^ × 60 μm^2^ to 5 μm^2^ × 5 μm^2^, and the scanning frequency is 1 Hz. The mechanical measurement of single-cell was detected using contact mode in RPMI-1640 medium at 1.63 m/s of moving forward speed and 0.8 Hz of scanning speed. DNP-S of AFM probe was used to collect the curve of force-distance, and the resonant frequency of probe ranged from 12 kHz to 24 kHz, the force elastic coefficient of microcantilever was 0.06 N/m, the length or width of microcantilever was 205 μm or 25 μm respectively. At least 15 cells were selected and more than 1,000 curves of force-distance were measured in each group. Young's modulus and adhesion force of Gaussian fitting were analyzed and calculated using Nano Scope analysis software 8.14.

### Quantitative real-time polymerase chain reaction (qRT-PCR)

The total RNA of JEC cells treated with 0.1 % DMSO or different concentrations of HNPG (2, 4 and 8 μM) for 24 h was extracted using Trizol (Invitrogen Inc., Carlsbad, California, USA). 1 μg of total RNA was reversely transcribed to cDNA using the Revert Aid™ Fisrt Strand cDNA Synthesis Kit (Fermentas Inc., Shanghai, Chian) according to manufacture protocols. The qRT-PCR procedure was performed using the SYBR^®^ Premix Ex Taq™ II (Tli RNaseH Plus) system (TaKaRa Inc., Dalian, Liaoning, China) in an Applied Biosystems 7500 Fast real-time PCR machine (ABI Inc., Carlsbad, CA, US). The primer sequences are shown in Table 1. The qRT-PCR program was set as follow initial denaturation at 95 °C for 5 min, 40 reaction cycles, with each cycle consisting of a denaturation at 95 °C for 15 s, annealing at 60 °C for 30 s, and then elongation at 72 °C for 30 s. GAPDH mRNA served as the internal control. The value of cycle threshold (CT) were recorded, and the mRNA levels of target genes was obtained using the following equation of ΔΔCT = (CT_purpose gene_ - CT_internal gene_)_treatment group_ - (CT_purpose gene_ - CT_internal gene_)_control group_ via computing the value of 2^-ΔΔCT^ of each group.

**Table. 1.**
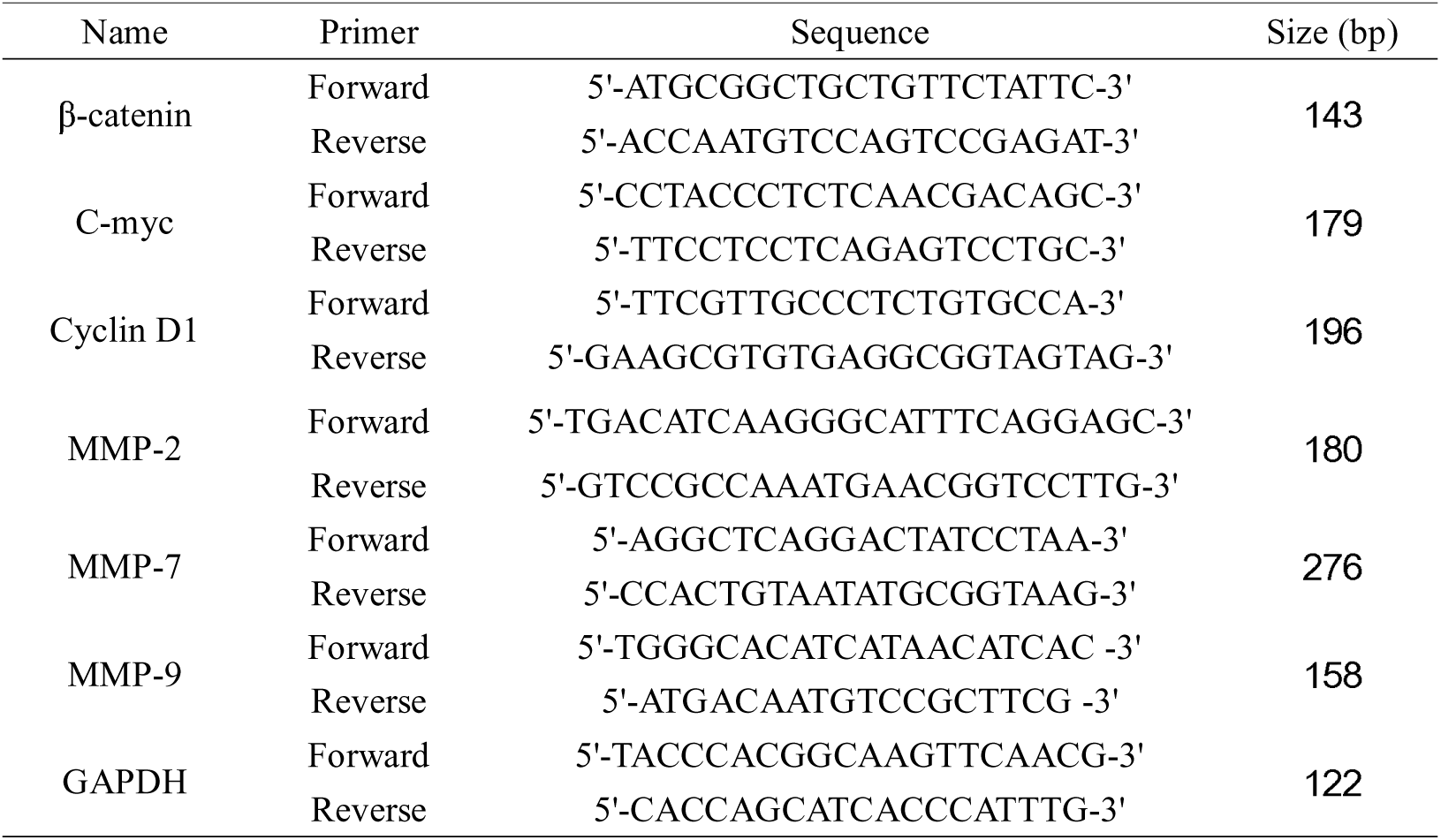
Primers for quantitative real-time polymerase chain reaction (qRT-PCR)

### Western blotting analysis

JEC cells that were administrated to 0.1 % DMSO or different concentrations of HNPG (2, 4 and 8 μM) for 24 h were lysed in 5 % lysis buffer (Tiangen Biotech Co., Ltd., Beijing, China). The amount of total protein was determined by BCA kit (Tiangen Biotech Co., Ltd., Beijing, China). Protein (20 μg) was separated by 10 % SDS-PAGE and transferred to nitrocellulose membranes. Non-specific binding sites were blocked by incubating the nitrocellulose membrane for 1 h at 37 °C with 5 % non-fat dried milk in TBST. The membranes were incubated for overnight at 4 °C with primary antibodies (rabbit anti-β-catenin, rabbit anti-C-myc, rabbit anti-Cyclin D1, rabbit anti-MMP-2, rabbit anti-MMP-7, rabbit anti-MMP-9 and rabbit anti-GAPDH) on shaker, then a horseradish peroxidase-conjugated secondary rabbit antibody for 1 h at 20 °C on shaker. Bands were visualized using an enhanced chemiluminescence system (Catalog Number: AR1170; Boster Biological Technology Co., Ltd., Wuhan, Hubei, China) and analyzed using Alpha Image 2200 (Version: 1.0; National Institutes of Health, Maryland, US). Each experiment was repeated three times and the mean value was obtained.

### Statistical analysis

SPSS 18.0 software package (SPSS Inc., Chicago, IL, USA) was used for analysis. Data are presented as the mean ± standard deviation. The means of multiple groups were compared with one-way analysis of variance (ANOVA), P<0.05 was considered to indicate a statistically significant difference.

## Acknowledgements

The authors would like to thank Professor Manpeng Huo (the Medical School of Yanan University, Yanan, Shannxi, China) for his technical assistance.

## Author contributions

J. B. designed and performed experiments, analyzed and explained data, wrote, edited and reviewed manuscript. B. Y. performed experiments, analyzed data. XL: administrated funding, designed research, edited and reviewed manuscript.

